# Exploring the Potential of Large Language Models in Molecular Tasks: An Insightful Evaluation with GPT‐4

**DOI:** 10.1101/2023.11.28.568966

**Authors:** Jinlu Zhang, Yin Fang, Ningyu Zhang, Xin Shao, Huajun Chen, Xiaohui Fan

## Abstract

In the rapidly changing realm of artificial intelligence, large language models (LLMs) such as GPT-4 are increasingly being explored for their potential to aid and enhance the field of molecular research. This study explores the performance of GPT-4 and GPT-3.5 in molecular research, particularly in generating and optimizing molecular structures. The results highlight GPT-4’s strengths in certain areas of molecular optimization, while also revealing challenges in accurately generating complex molecules. The findings underscore the necessity for integrating these models with domain-specific tools to enhance their application in scientific research, particularly in molecular studies. The study offers insights into the potential of LLMs for advancing molecular research, paving the way for future developments in this rapidly evolving field.

## 1 Introduction

Molecular research plays an important role in understanding life processes, disease mechanisms, and designing drugs[1]. This field’s significance stems from its direct impact on healthcare and medicine, providing insights into the building blocks of life and potential pathways for therapeutic interventions. However, traditional methods in molecular research are often challenged by the complexity of molecular structures, necessitating substantial time and resources[2][3][4]. Such complexity can affect the pace of scientific endeavors, especially in vital areas like drug discovery where rapid development and innovation play key roles[5][6].

In this context, the rapid evolution of large language models (LLMs) has emerged as a potential game-changer[7]. These models, particularly GPT-4[8] from OpenAI, are redefining the boundaries of artificial intelligence with their advanced capabilities in language processing and understanding. Their development marks a significant shift not just in computing but in how various fields, including molecular research, can leverage AI. By tightly integrating language and science, these models initiate a new approach based on language in chemistry and biology research[9][10][11]. For instance, in drug design, the use of large language models can lead to more efficient simulation of molecular design and optimization processes on computational platforms. This integration has the potential to significantly shorten the drug development cycle and reduce associated research and development costs[12][13][14].

To explore the practical impact of LLMs in molecular research, our study focuses on evaluating GPT-4 and GPT-3.5 across three specific molecular tasks: molecular description generation, molecular optimization, and molecular generation. We investigate whether these AI models can effectively generate accurate molecular descriptions from SMILES[15][16] data, optimize molecular structures to enhance drug-likeness, and create molecules in line with provided descriptions. This evaluation seeks to understand the capacity of these primarily language-based AI models in addressing the complexities inherent in molecular research.

The findings from our research illuminate the capabilities and limitations of LLMs in molecular science, contributing to the broader conversation on AI’s role in scientific discovery. Our analysis emphasizes the need to integrate these models with external tools and validation techniques, such as computational chemistry and wet lab experiments, to ensure their practicality and accuracy. This collaborative approach between AI and human expertise in molecular research heralds a future of enhanced innovation, where AI augments the human capacity for discovery and development in healthcare and medicine. A summary of our findings and their implications can be found in Figure 1.

**Fig. 1:**
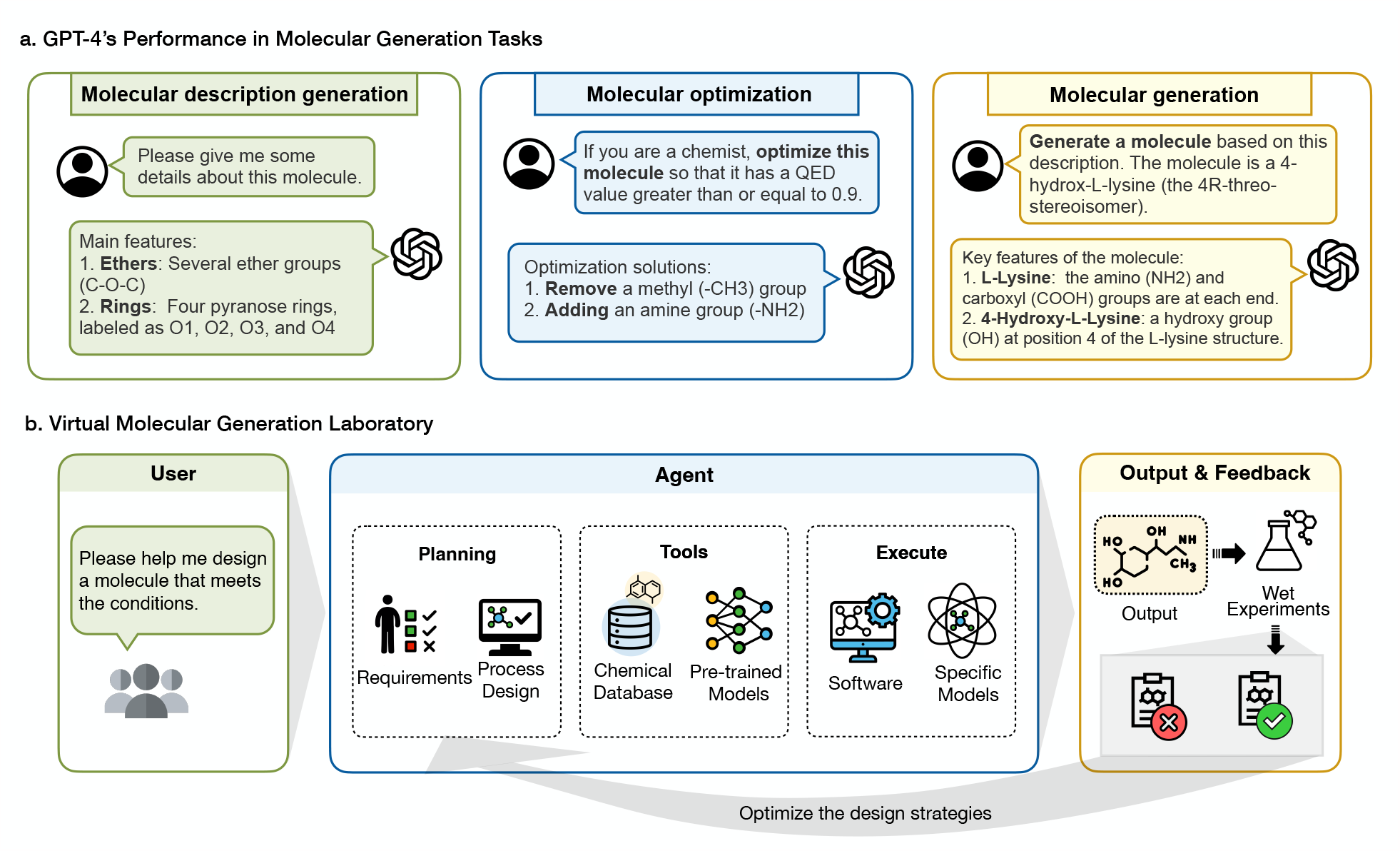
GPT-4’s performance in molecular generation tasks and virtual molecular generation labs.

## 2 Results

### Molecular description generation

Molecular description generation is a very important step in the fields of molecular design, drug discovery, etc. It can help chemists and biologists to quickly obtain information about the molecule and guide experiments and research directions.

As illustrated in Figure 2, we can note that both GPT-3.5 and GPT-4 do not answer the question directly, but rather break the molecule down into various different functional groups based on the structure of the molecule and describe in turn the role that each functional group may play in the molecule. The correct answer to this case is “The molecule is a steroid ester that is methyl (17E)-pregna-4,17-dien-21-oate substituted by oxo groups at positions 3 and 11. positions 3 and 11. It is a 3-oxo-Delta(4) steroid, an 11-oxo steroid, a steroid ester and a methyl ester. It derives from a hydride of a pregnan. The molecule is an ester, and both GPT3.5 and GPT-4 profiled his functional group, but GPT-4 was able to point out that it had an ester group, while GPT3.5 failed to do so. This indicates that GPT-4 has a better understanding of the molecular structure and functional groups compared to GPT-3.5. Moreover, this case illustrates that although GPT-4 only gives molecular descriptors and cannot directly give information such as specific names, properties or potential applications of molecules, its analysis of molecular functional group splitting is essentially correct.

**Fig. 2:**
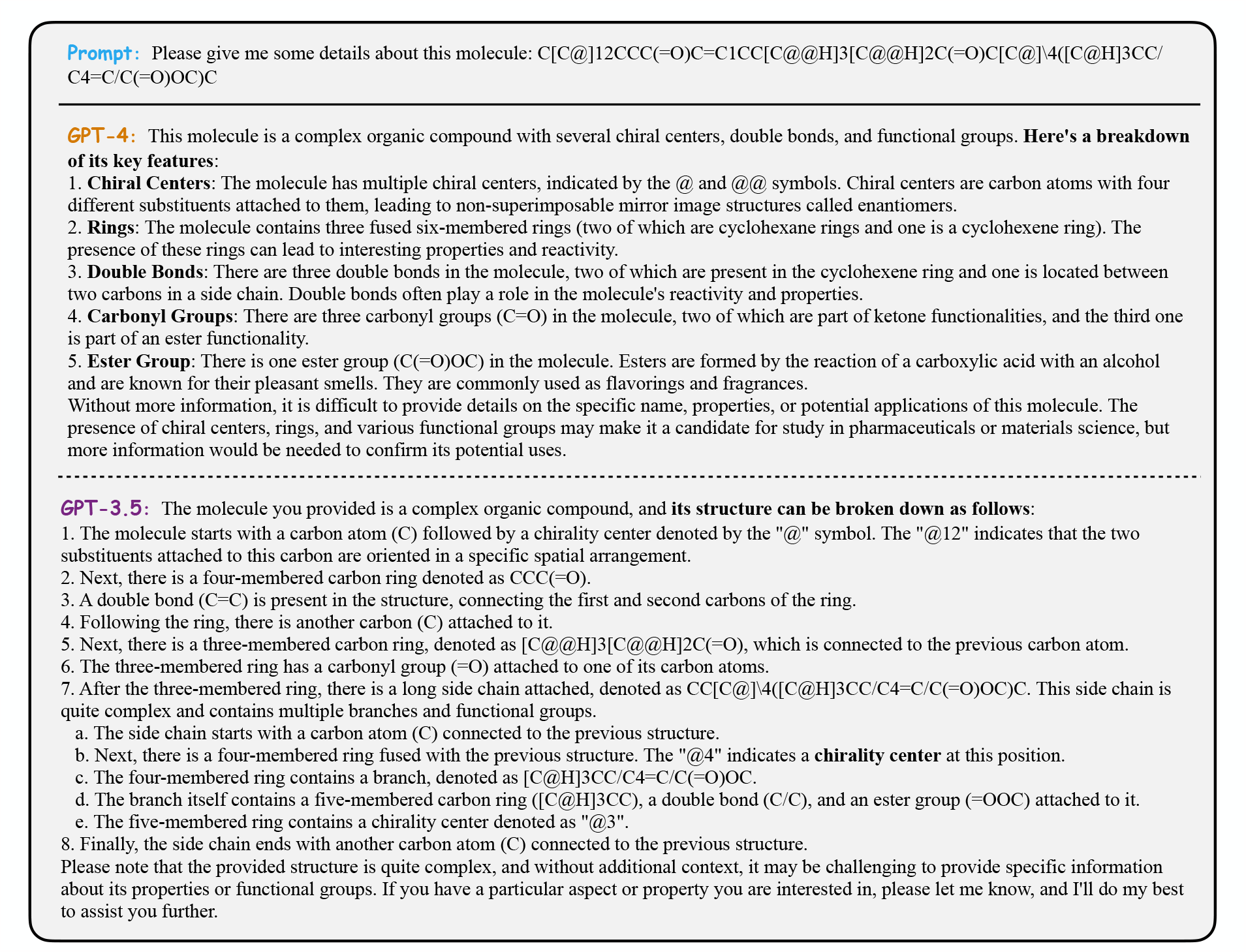
Molecular Description Generation Test Result.

### Molecular optimization

Molecular optimization can be used to study the relationship between the structure and properties of molecules, to optimise the stability, reactivity, adsorption and other physical properties of molecules, and it can also be applied in the fields of drug design, catalyst design and material design[17].

In the test, we asked the same questions for GPT-3.5 and GPT-4. GPT-4 proposes different optimization schemes for different molecules and gives the corresponding smiles expressions. In contrast, the optimization schemes proposed by GPT-3.5 for different molecules are all the same and and refuses to give the corresponding SMILES. We counted the answers given by GPT-4 and calculated the QED values using the functions in the rdkit package. In more cases of optimization of QED values of molecules, we can find that although GPT-4 sometimes generates some invalid molecules, it still proposes quite a few strategies for molecules that give better QED values. GPT-4 is able to propose specific modification steps for a given molecule and provide the modified molecule. However, these molecules do not necessarily fulfill the predefined requirements. In the test case shown in Figure 3, we calculated the QED values corresponding to the four modified molecules given by GPT-4. We found that only one of the four answers given by GPT-4 has a higher QED value than the original molecule, but it failed to reach 0.9 and some of the SMILES expressions given were invalid. With the optimized and modified molecules by GPT-4, the QED value was successfully improved in 48.39% of cases. Moreover, during the optimization process, GPT-3.5 and GPT-4 only accept inputs in the form of SMILES, and for all other molecular descriptors they were unable to give an answer. This may be related to its pre-training corpus.

**Fig. 3:**
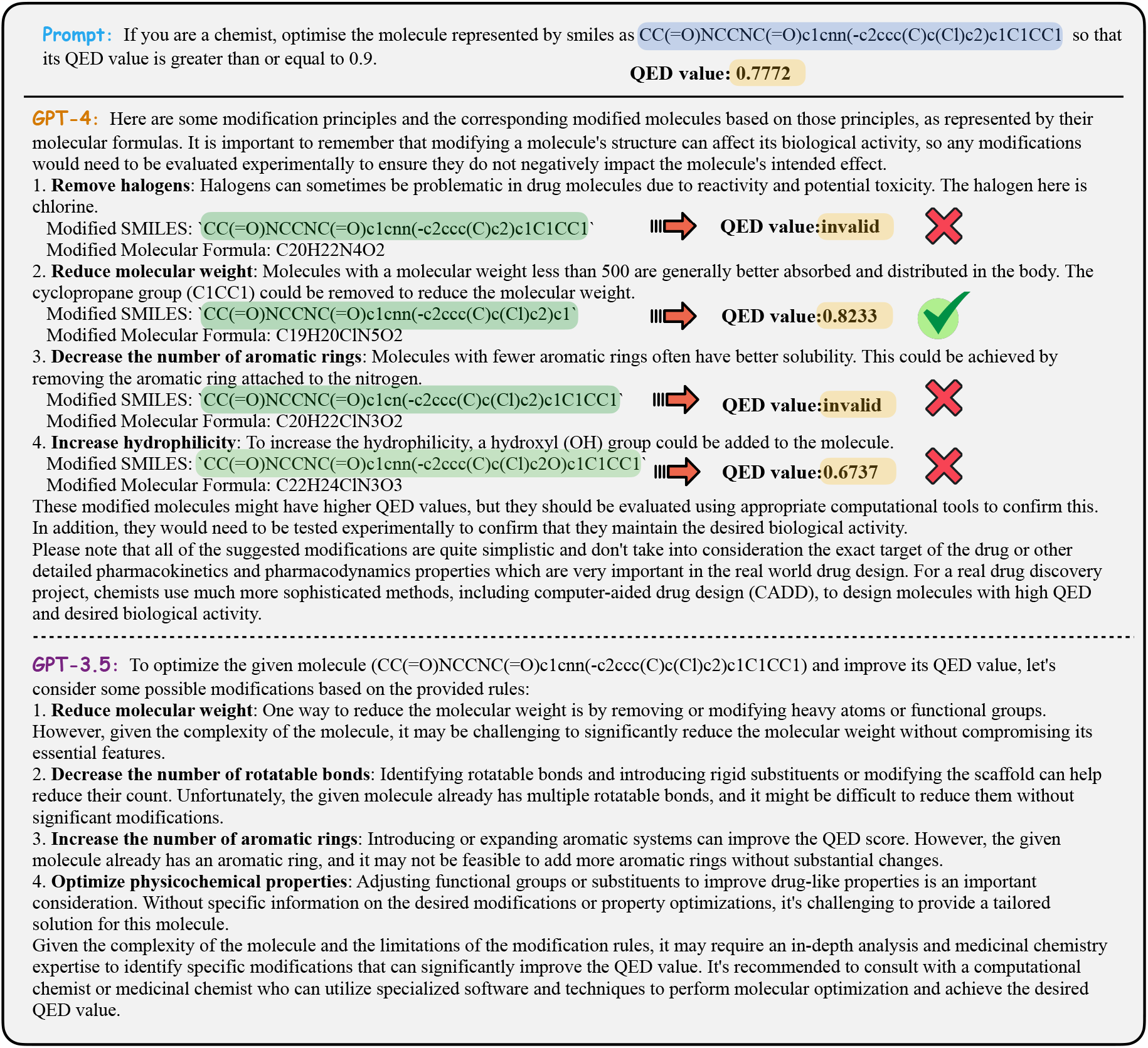
Molecular optimization Test Result.

### Molecular generation

In addition to molecular description generation, whether GPT3.5 and GPT-4 can generate a compliant molecule based on a piece of molecular description is also one of our concerns. Description-based molecule generation can help drug researchers to rapidly generate molecules with specific structures and properties for drug screening and design.

Both GPT-3.5 and GPT-4 give a possible SMILES descriptor and summarize some features of the molecule. However, GPT-4 has some errors in the description process, as shown in the red part of Figure 4, in which there is no carbon-carbon double bond between C17 and C18. For a more intuitive comparison, we visualized the SMILES descriptors generated by GPT-3.5 and GPT-4 as molecular diagrams. As shown in Figure 4, the molecules on the blue background are the molecules corresponding to the text of this descriptor (i.e., the input text describes the structure and properties of the molecule to its right), and the molecules at the bottom are the molecules generated by GPT-4 and GPT-3.5 based on this descriptor. It can be seen that 1) the molecules visualized according to the SMILES descriptor do not correspond to the structures of the molecules provided above, and 2) the two molecules generated by both have a certain degree of similarity to the structures of the real molecules. Although all three are steroid compounds, the generated molecules do not exactly match the descriptor text.

**Fig. 4:**
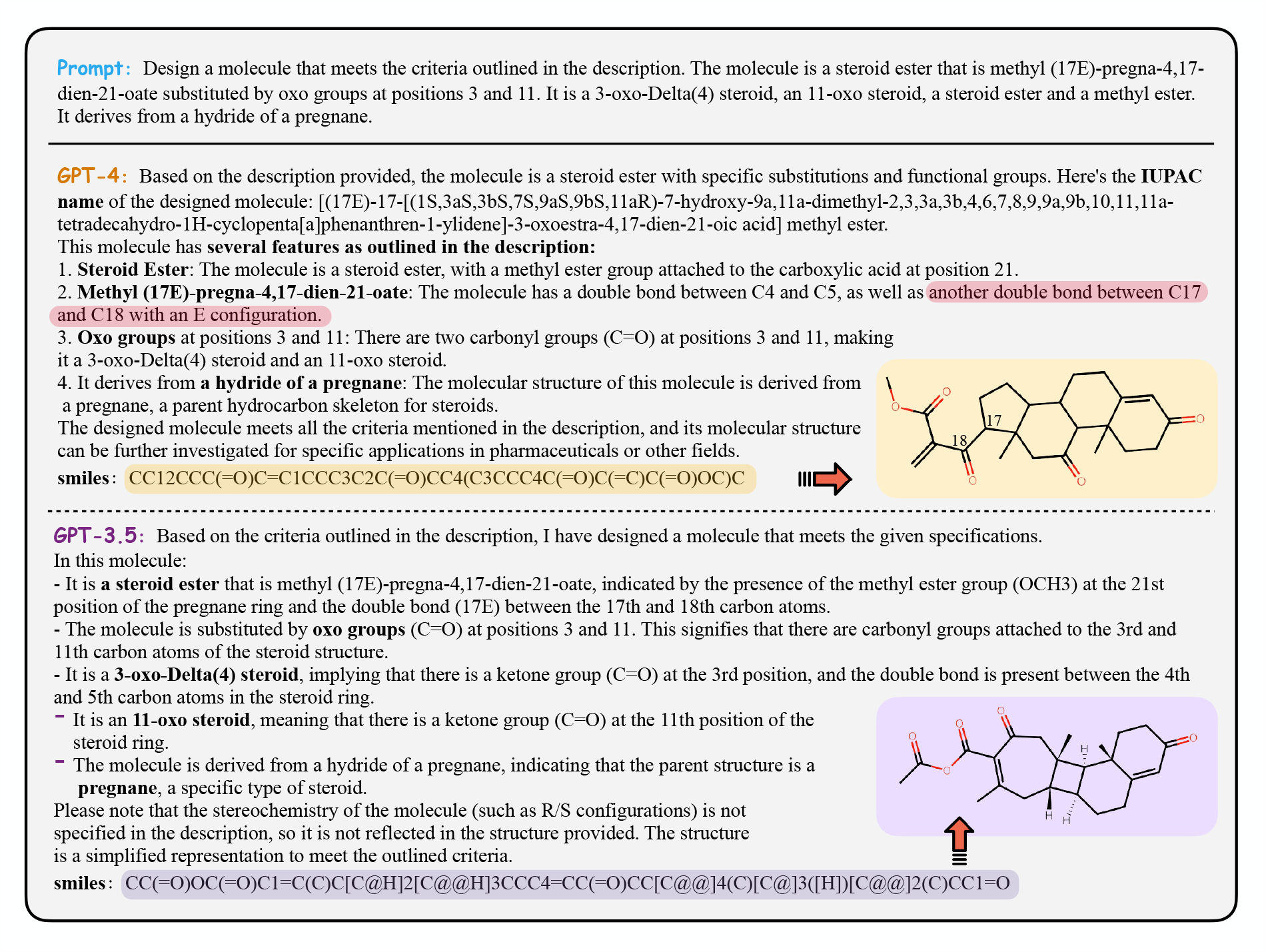
Molecular generation Test Result (The part in red indicates an error, there is no carbon-carbon double bonds in the provided SMILES.)

The results showed that GPT-4 outperformed GPT-3.5 in all three tasks. In the task of generating molecular descriptions, GPT-4 outperforms GPT-3.5 in its understanding and depiction of functional groups. However, when it comes to molecular modification, GPT-3.5’s capabilities are limited to providing suggestions, as it lacks the ability to deliver the modified molecule. On the contrary, GPT-4 demonstrates a remarkable proficiency, with a success rate as high as 48.39% in modifying molecules to achieve a desired Quantitative Estimate of Drug-likeness (QED) value. In the realm of molecular generation, both GPT-3.5 and GPT-4 can generate some molecules, but they often fall short of fully satisfying the provided description. Only some functional groups could be resolved, and the overall success rate stayed at 0%. This indicates that these models still face challenges in understanding complex molecular descriptions and translating them into specific molecules. It is likely because these large language models are trained to learn only a fraction of text and are not specifically designed to handle the complexities of molecular structure and behavior.

## 3 Discussion

The performance of GPT-4 and GPT-3.5 in our molecular research tasks highlighted both their potential and limitations. While GPT-4 demonstrated enhanced capabilities, particularly in molecular optimization, both models struggled with the accurate generation of complex molecules. This suggests that despite their advanced capabilities, LLMs like GPT-4 and GPT-3.5 still require further refinement to fully meet the demands of molecular research.

Currently, there are relatively few large language models developed specifically for the molecular domain. This is due to the unique structures and features of molecules, which demand specific optimization goals and training methods for developing large language models capable of comprehending and generating molecular information more effectively. By defining precise optimization goals, the large language model can be guided to acquire the structural information and behavioral patterns of molecules with greater accuracy. This process may encompass the utilization of particular loss functions, data pre-processing methods, or enhancements in model structure in order to facilitate the improved capture of the molecule’s specific properties. In the case of drug molecule design, for example, this process involves optimising the pharmacokinetic (PK) and pharmacodynamic (PD)[18] properties of a drug. If a large language model can deeply analyse and learn the structural, stereochemical and physicochemical properties of a compound, then it will be possible to predict processes such as the distribution, metabolism and excretion of a drug in the body. GPT-4’s multimodal processing capabilities and the new Assistants API allow it to be integrated with specialised chemistry tools and systems to improve prediction accuracy. This predictive capability is extremely important as it relates to the bioavailability, selectivity and safety of the drug. In this way, we can design more effective drug candidate molecules. Another avenue of research involves integrating multiple agents into an intelligent system[19]. For instance, in a complex task of generating molecules, various specialized agents can collaborate and invoke suitable large language models to collectively work together. Some agents may be responsible for parsing user instructions and planning, while others may focus on information retrieval, molecular structure generation, or verification of the correctness of the resulting molecules, among other responsibilities. This collaborative working model allows the whole system to be more flexible and efficient, and can be used for self-designed experiments to generate new drug molecules, for example, to better meet the needs of users.

Additionally, validation is a crucial challenge for ensuring the accuracy and credibility of the generated molecules. Although large language models have the ability to generate molecules, external tools and wet experiments are required to validate and verify the accuracy of the results. Computational chemistry tools[20] are highly valuable in this context as they enable molecular simulations and predictions, offering insights into the physical and chemical properties of molecules, including stability and reactivity. Comparison with experimental results serves as a means of verifying whether the generated molecules meet expectations. If there are discrepancies between predictions, experimental results, and expectations, these can be used to optimize and adjust the model via the feedback mechanisms inherent in artificial intelligence and machine learning. This approach enhances the model’s predictive accuracy and generative abilities. For those molecules deemed valuable for research or potential applications, it is essential to carry out empirical testing through wet experiments in the field. However, the choice of suitable experimental validation methods and techniques remains a significant challenge. Techniques such as nuclear magnetic resonance (NMR) and mass spectrometry (MS) help determine molecular structure, while infrared (IR) and ultraviolet-visible (UV-Vis) spectroscopy probe optical properties. By synthesizing these molecules and subjecting them to experimental testing, it becomes possible to validate the model’s generation accuracy and credibility against actual properties. This, in turn, bolsters confidence in the model’s ability to generate viable molecular structures.

## 4 Method

### 4.1 GPT-3.5 and GPT-4 versions

The GPT-3.5 version used was between the dates 4/01 to 4/12/2023. The GPT-4 version was between 4/10 to 4/23/2023.

### 4.2 Datasets

We used the dataset ChEBI-20[21] for the molecular description generation and molecular generation tasks, and the dataset ZINC[22] for the molecular optimization task.

### 4.3 Evaluation and Sampling

Our evaluation of GPT-4 and GPT-3.5’s capabilities in the molecular domain was conducted via ChatGPT’s interactive interface, using a random sampling approach. We chose 10 distinct cases for each task to ensure a comprehensive assessment. This methodology was designed to explore the generative and predictive capacities of the models across diverse molecular tasks.

It is important to note that our findings are based on randomly sampled data and interactive testing. This approach, while offering valuable insights, may be influenced by the specificities of data distribution and sample selection.

## 5 Data availability

All experimental data are available in Figshare (https://doi.org/10.6084/m9.figshare.23816079).

